# Spatial and temporal signatures of cell competition revealed by K-function analysis

**DOI:** 10.1101/2024.10.29.620688

**Authors:** Nathan Day, Jasmine Michalowska, Manasi Kelkar, Giulia Vallardi, Guillaume Charras, Alan R. Lowe

**Author notes:** These authors contributed equally (Nathan Day, Jasmine Michalowska).

## Abstract

It is generally hypothesised that in MDCK wild-type versus Scr^KD^ co-cultures mechanical competition leads to the physical elimination of Scr^KD^ cells, while in MDCK wild-type versus Ras^V12^ co-cultures biochemical competition results in the non-apoptotic extrusion of Ras^V12^ cells. However, the exact spatiotemporal signatures of competitive interactions in these systems remains largely unknown. Utilising advanced single-cell image analysis, this research quantifies the spatiotemporal dynamics of these competitive interactions. Time-lapse microscopy, combined with cellular segmentation, tracking, cell-state classification, and K-function clustering analysis provide detailed insights into how wild-type cell proliferation relates to the elimination of mutant cells. We find striking differences in the underlying processes of each competition type. In the Scr^KD^ competition, elimination seems driven by diffuse proliferative forces unrelated to the immediate elimination site. In contrast, Ras^V12^ cell elimination is closely linked to local increases in wild-type cell mitoses prior to the Ras^V12^ extrusion. Both competitions trigger an increase in local wild-type cell proliferation post-elimination, although the timing of these responses vary. This study not only sheds light on the diverse mechanisms of cell competition but also underscores the complexity of cellular interactions in tissue dynamics, providing new perspectives on cellular quality control and a new quantitative approach to characterise these interactions.

## 1 Introduction

Even simple cellular systems display incredible diversity; individual cells within populations of either clonal or closely related origin often exhibit highly heterogeneous behaviour as they respond to spatiotemporally varying environmental cues. One example of this complexity, where individual cell fate is determined by the local tissue organisation over time, is cell competition.

Cell competition was first discovered in *Drosophila melanogaster* Minute mutants [1]. A spatial dependency was later revealed, showing that mutant cell elimination was expedited by fragmentation of the mutant population [2]. These observations were the first that depicted a key aspect of this context dependency: otherwise viable cells are out-competed when placed within a mixed population and, subsequently, are eliminated [3, 4].

The majority of cell competition studies have categorised context dependent cell elimination into two broad categories; mechanical and biochemical, as shown in **Fig. 1**. Biochemical competition is defined as operating via mechanisms of cellular signaling between cells in contact with one another [5, 6]. Here, factors such as cell identity recognition [7, 8] and competition for resources like growth factors are important in driving outcomes. MDCK wild-type co-cultured with mutant cells over-expressing oncogenic Ras (Ras^V12^) has been shown to operate via a context dependency that is suggestive of cell-cell interfacial recognition, a key hallmark of biochemical competition. The competitive elimination in this scenario is also understood to depend upon the apical extrusion of the mutant Ras^V12^ population [9]. Mechanical competition is defined by a cell’s response to physical stresses [10], including compression, strain, cell migration, mitotic forces and tissue relaxation due to apoptosis or extrusion [11, 12]. MDCK wild-type populations co-cultured with mutant cells depleted in the polarity protein *scribble* (Scr^KD^) [13] has previously been shown to exhibit mechanical competition [12] where the mutant cells are eliminated due to a hypersensitivity to compressive strain.

**Fig. 1.**
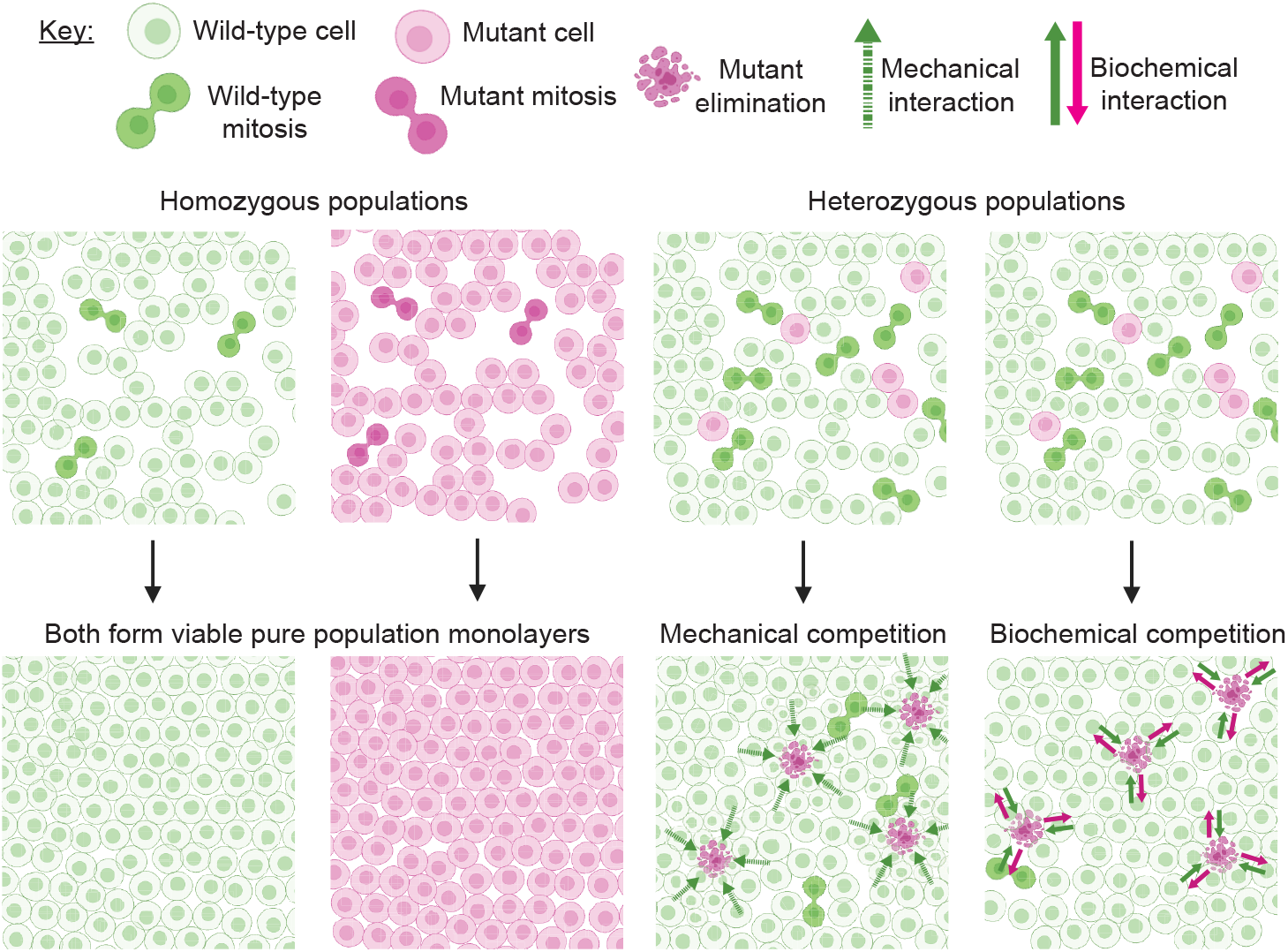
A comparative illustration of the canonical definitions of biochemical and mechanical cell competition. Cell competition is initiated when mixing a wild-type population (green cells) with a mutant population (magenta cells). Both cell types are viable as pure populations and it is only upon mixing that they exhibit competitive elimination behaviour. Competitive elimination has typically been characterised as either mechanical, with competitive insults traded over longer distances via compression and strain of the monolayer, or biochemical, where interactions between the two populations occur between nearest neighbours via mechanisms of cellular signaling. [30]

Importantly, there is nothing to suggest that the two categories of biochemical and mechanical competition are mutually exclusive. Ras^V12^ cell competition is understood to operate on a biochemical basis [14], but possesses an implicitly mechanical method of mutant elimination in the form of apical cell extrusion. On the other hand, in mechanical Scr^KD^ competition, the division of wild-type cells appears favoured in neighbourhoods with many mutant cells [15], something which is indicative of a biochemical type of cell interface recognition. The indistinct nature of these biological/mechanical categorisations, and the clear overlap of these categories when observing various types of competition, opens the space to a more nuanced model of cell competition.

To begin to quantitatively understand how dynamic, cell-scale topological, biochemical and physical parameters determine the precise mechanisms of cell fate research in the field has increasingly adopted in-silico models of competitive co-cultures [14, 16, 17]. We previously employed a Cellular Potts model to examine Scr^KD^ cell competition dynamics by integrating mechanical features such as cell-cell affinity, cell-substrate adhesion, cellular velocity and cell compressibility [14]. This particular study found that the two main parameters that contributed the most to the competitive outcome of Scr^KD^ elimination were found to be the difference in homeostatic density between the two populations and the relative compressibility of the cells. Further work from this lab, using quantitative, single-cell methods, has shown that in Scr^KD^ competitive environments, cell death is driven by an increased incidence of mitosis in the wild-type population [15]. However the exact details of the temporal causality in this system are still unknown and the question of whether the mutant cell elimination is a cause of consequence of a concurrent increase in wild-type cell density remains.

In this paper, we expand upon this work by pursuing an answer to the question of causality in competitive systems using novel single-cell quantitative methods. We combine the analysis of population-level dynamics in pure and mixed co-cultures with K-function cluster analysis to specifically understand the spatial and temporal dependency of the interplay between wild-type mitosis and mutant cell elimination events.

We propose that the locations and timings of these mitotic events is one key phenomenon that can then be used to quantitatively categorise competing cell populations into four distinct categories; localised versus diffuse, and causal versus induced competition. Categorisation of competition along these lines would be based on how far the effects of mitosis can be felt throughout the tissue: whether there is an immediate cell-interface type recognition and reaction to mitosis, or whether the reaction is instead more diffuse and felt as an increase in density and tissue-wide strain that eventually results in elimination. Further to this spatial consideration, the timing of competitive interactions is also crucially important in the trajectories of these competitive outcomes. If the local wild-type cellular environment displays atypical behaviours (be it concerted migration or growth dynamics) in the time period preceding the mutant cell elimination, then this is suggestive of a causative impact. If the wild-type behaviour deviates from the norm only after the mutant cell elimination then this is suggestive of a wild-type reaction to the competition, rather than an instigation of it.

The analytical methods and classification framework of competition presented here can be generally applied to quantify the spatiotemporal dependency of events in competitive co-cultures and to understand cell interactions with relevance to both cancer and stem cell biology.

### 1.1 Experimental and computational strategy

To characterise the spatial and temporal signatures of cell competition a highthroughput imaging-to-analysis pipeline was developed. Time lapse fluorescence microscopy was performed over multiple days, imaging co-cultures of MDCK^WT^cells expressing the H2b–green fluorescent protein (H2b-GFP) vs Ras^V12^ mutants expressing the H2b–red fluorescent protein (H2b-RFP), in the ratios 99:1, 97:3 and 95:5 wild-type:Ras^V12^, respectively. MDCK^WT^cells vs Scr^KD^ mutants expressing the H2b–red fluorescent protein (H2b-RFP) are imaged in the ratio 90:10 wildtype:Scr^KD^, respectively. Pure population MDCK^WT^, Ras^V12^ and Scr^KD^ populations are also imaged in the same way.

A bespoke automated epifluorescence microscope built inside a standard CO_2_ incubator is used to acquire brightfield, RFP and GFP images every 4 minutes for each population type with a 20x air objective. 717 × 558 μm^2^ regions are acquired for 1200–1400 frames using 24-well imaging plates. Competitions and controls are run simultaneously in identical conditions and each condition is run *n*⩾ 3 independent times. Pre-processing steps include cropping all frames to 640 x 480 μm^2^ regions containing 800–1000 cells per field of view.

Cell nuclei are segmented using *Cellpose*[18], *StarDist* [19, 20] or a residual U-Net [21]. Each cell instance was tracked over time using *btrack* [15, 22]. An “event” refers to either a cell mitosis, apoptosis or extrusion, either identified manually or by using a dedicated convolutional neural network previously described [15, 22, 23]. For the purpose of the population-level analysis all apoptosis and extrusion events are referred to as ‘elimination events’. Here where both extrusion and apoptosis events occurred for a particular cell population they are combined. Examples of mutant elimination events can be seen in **Fig. 2**. Ras^V12^ extrusion events are defined as the time point at which the cell has rounded up and moved out of the focal plane and are manually identified. Apoptotic and mitotic cell state transitions are identified as part of the tracking pipeline with the precise time point of each apoptosis event manually verified to enhance precision. In competitive co-cultures a total of *N*_WT mitosis_ = 14, 919 events, *N*_Ras elimination_ = 90 events and *N*_Scr elimination_ = 101 events were identified.

**Fig. 2.**
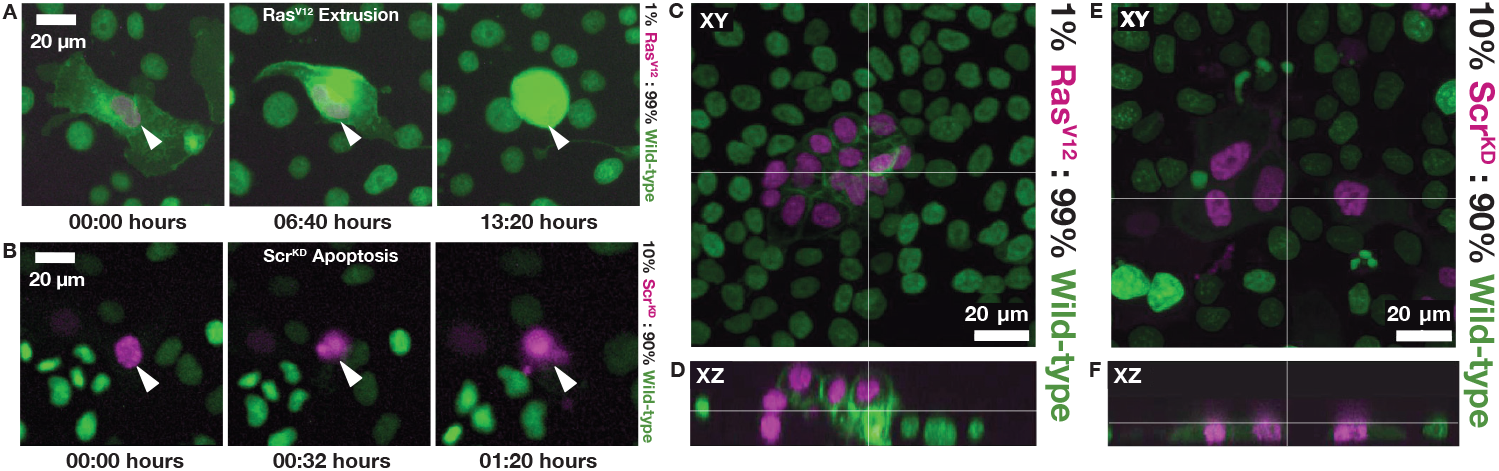
Examples of mutant elimination events in cell competition. Mutant cells (Ras^V12^ and Scr^KD^) are shown with magenta nuclei (H2B-RFP) and wild-type cells are shown with green nuclei (H2B-GFP). **(A)** Live-cell image sequence showing the temporal progression of Ras^V12^ apical extrusion. **(B)** Live-cell image sequence showing the temporal progression of Scr^KD^ apoptosis. **(C, D)** Confocal z-stack images depicting fixed apical extrusion of Ras^V12^ cells in the context of wild-type cells. **(E, F)** Comparative confocal z-stack images depicting fixed Scr^KD^ cells in the context of wild-type cells, with no apical extrusion. There is a 1μm step between z-slices.

## 2 Results

### 2.1 Quantification of population-level dynamics in pure and mixed populations reveals distinct differences between Ras^V12^ and Scr^KD^ competition

#### 2.1.1 Comparative analysis of pure and mixed co-cultures displays differing growth rates between wild-type populations in Ras^V12^ and Scr^KD^ competition

Following cell segmentation, tracking and event identification the growth dynamics of both pure and competitive co-cultures were quantified on a population-scale to confirm consistency with previous findings [15], and to identify any clear differences that may be present between Ras^V12^ and Scr^KD^ competition. To visualise cell population growth the per-frame cell count is extracted for *n* ⩾ 3 experiments per condition. The resulting population growth curves are displayed in **Fig. 3** alongside representative experimental images.

**Fig. 3.**
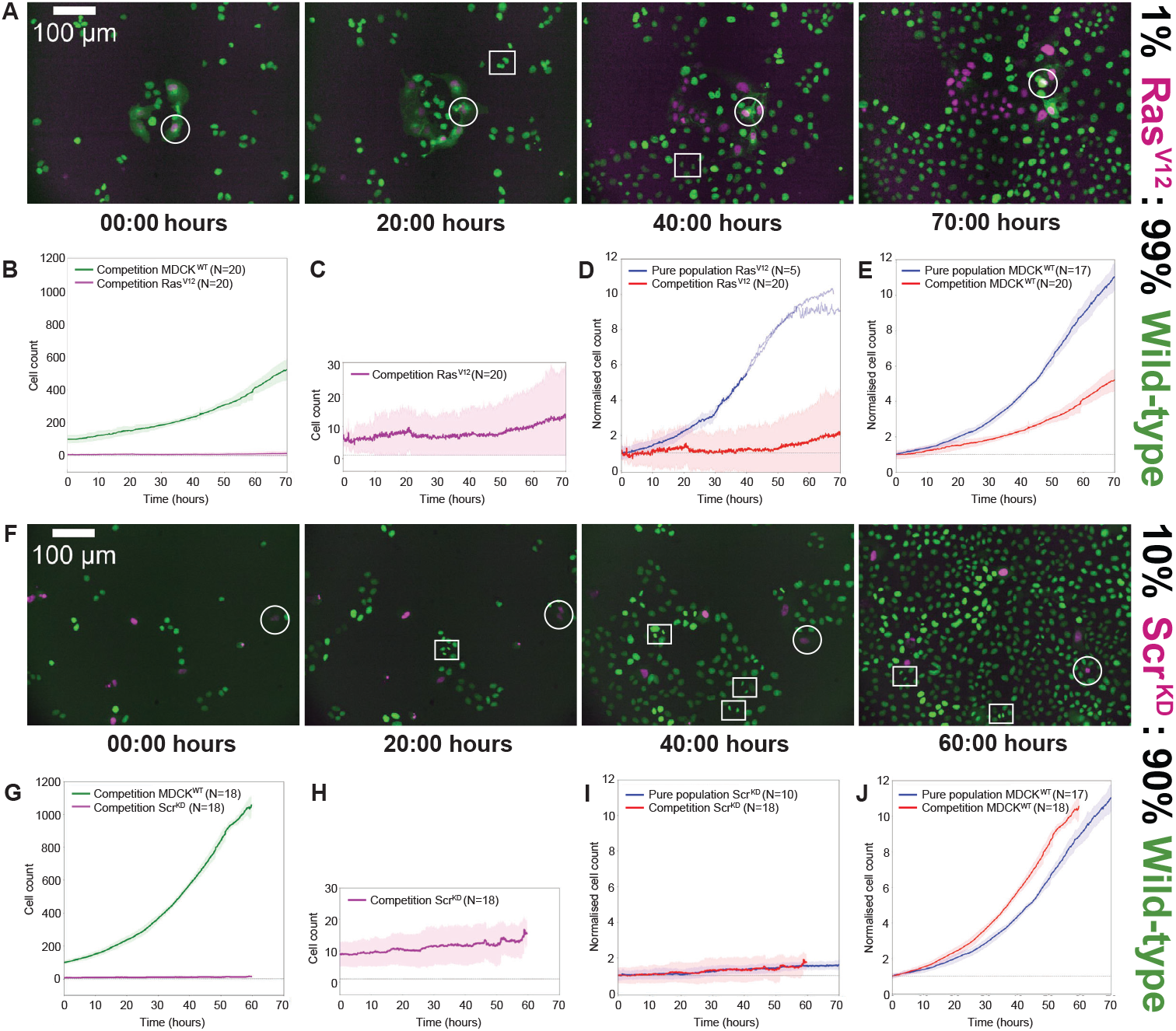
Cell population growth in pure and mixed ratios for two forms of cell competition. **(A)** Time lapse image sequence of competition between MDCK^WT^and Ras^V12^ cells in an initial 99:1 ratio. A single Ras^V12^ cell is followed across time with a white circle, ending in apical extrusion. MDCK^WT^mitosis events are highlighted with white squares. **(B)** Growth curves showing the suppression of Ras^V12^ growth in the presence of MDCK^WT^(wild-type) proliferation. **(C)** Magnification of the growth profile of the Ras^V12^ population. **(D)** Normalised growth curves showing the difference in proliferation for pure populations of induced Ras^V12^ versus Ras^V12^ populations in competition. **(E)** Normalised growth curves showing the difference in proliferation for pure populations of wild-type versus wild-type populations in Ras^V12^ competition. **(F)** Time lapse image sequence of competition between wild-type and Scr^KD^ cells in an initial 90:10 ratio. A single Scr^KD^ cell is followed across time with a white circle, ending in apoptosis. MDCK^WT^mitosis events are highlighted with white squares. **(G)** Growth curves showing the suppression of Scr^KD^ growth in the presence of MDCK^WT^(wild-type) proliferation. **(H)** Magnification of the growth profile of the Scr^KD^ population. **(I)** Normalised growth curves showing the similarly slow growth for pure populations of induced Scr^KD^ versus populations in Scr^KD^ competition. **(J)** Normalised growth curves showing the difference in proliferation for pure populations of wild-type versus wild-type populations in Scr^KD^ competition.

The Ras^V12^ versus wild-type cell counts displayed in **Fig. 3B** clearly display two populations with significantly different growth profiles over the course of the time-lapse acquisition. The wild-type population have a faster average growth rate than the Ras^V12^ cells, eventually reaching a cell count over 50 times greater than their mutant counterparts. This difference is partial evidence of competition, and to confirm this is not solely a consequence of the 99% wild-type to 1% Ras^V12^ initial seeding ratio the population dynamics of mixed co-cultures are compared to Ras^V12^ and MDCK^WT^grown in non-competitive, pure populations. Observing these in **Fig. 3D** and **Fig. 3E** respectively, a higher growth rate is observed in both of the non-competitive populations. Here, the pure Ras^V12^ population has a 5-fold higher growth rate compared to mixed population Ras^V12^ whereas the pure wildtype a 3-fold higher growth rate compared to mixed population wild-type. These differences indicate a suppression of growth rates in co-culture and suggests that there is an interplay between the Ras^V12^ and wild-type cells that impacts upon both population’s capacity to proliferate. The context dependency of the behaviour of both cell populations indicates that cellular competition is definitively taking place.

Differing growth rates between the Scr^KD^ versus wild-type populations are again clearly visible in **Fig. 3G**. Compared to Ras^V12^ competition the different cellular context of the Scr^KD^ cells appears to encourage the wild-type cells to proliferate at a faster rate, reaching a cell count number twice that of the wild-type cells in the Ras^V12^ competition in a comparatively shorter time frame. There is only a small difference when comparing the growth of non-competitive populations of Scr^KD^ against competitive populations, in what is likely the noticeable effects of Scr^KD^ ‘s hyper-sensitivity to cellular density discouraging the population against rapid proliferation in both scenarios [12]. In contrast to the Ras^V12^ competition, wild-type cells in Scr^KD^ competition grow faster than their non-competitive counterparts. This up-regulation of proliferation is consistent with prior results, where the probability of wild-type mitosis is higher in neighbourhoods populated by Scr^KD^ cells [15].

#### 2.1.2 Quantification of event dynamics in pure and mixed co-cultures indicates context-dependent regulation of mitotic behavior in wild-type, Scr^KD^ and Ras^V12^ populations

To further assess the population-level dynamics the number of automatically-classified mitoses and manually-identified elimination events were quantified. The Ras^V12^ versus wild-type cumulative mitosis events over time are shown in **Fig. 4A**. The corresponding Scr^KD^ versus wild-type cumulative mitosis curves are shown in **Fig. 4G**. Both of these graphs mirror the overall cell counts seen in **Fig. 3**, with wild-type mitoses greatly outnumbering both Ras^V12^ and Scr^KD^ mitoses. Comparing the two mixed co-cultures there is a much greater incidence of wild-type mitoses in the Scr^KD^ competition, explaining the larger population numbers when compared to the Ras^V12^ competition.

**Fig. 4.**
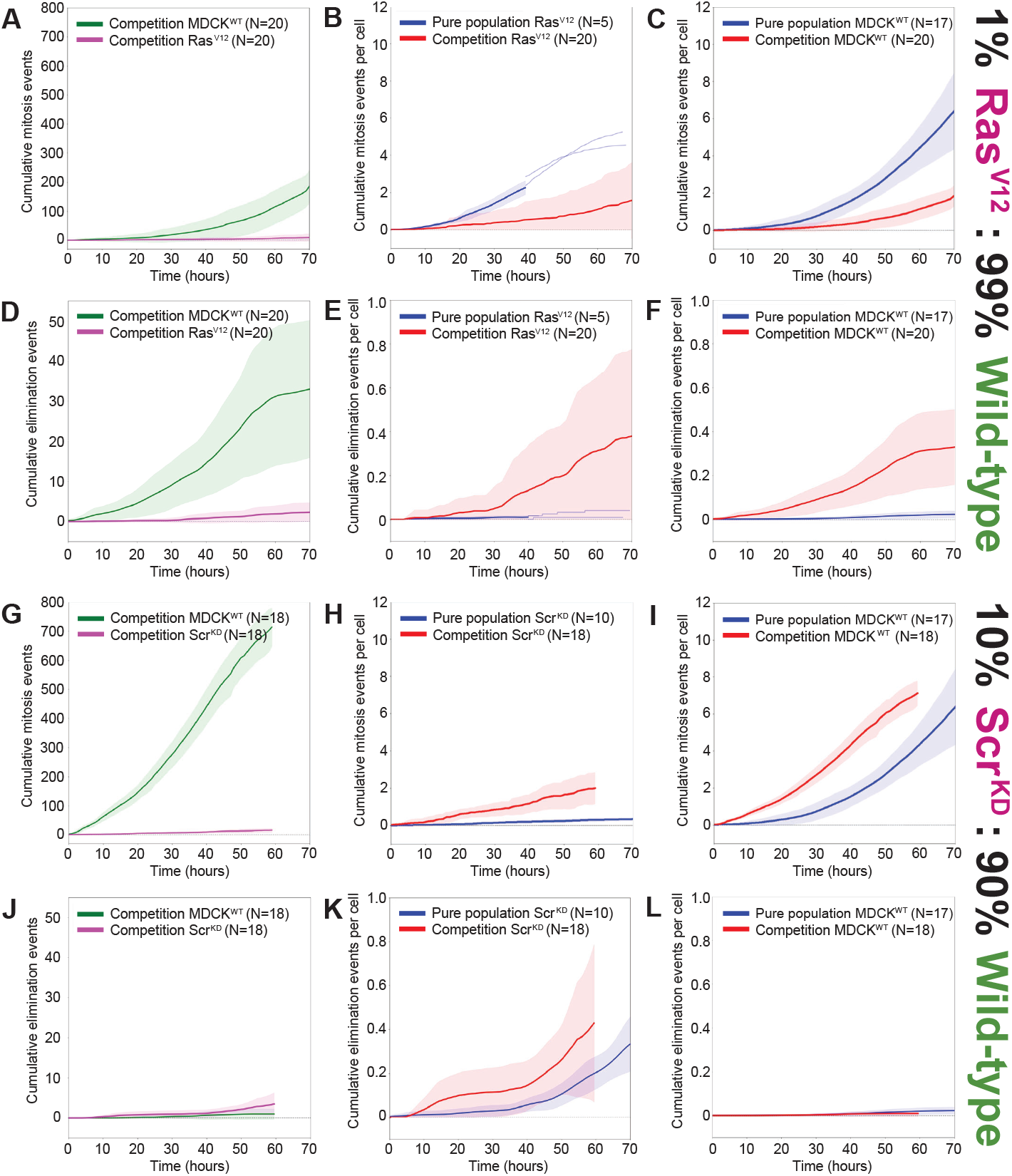
**Cumulative mitosis and elimination counts in pure (non-competitive) and mixed (competitive) conditions further differentiates the two forms of competition and raises the question of the causal relationship between mitosis and elimination**. **(A)** Cumulative mitosis curves for Ras^V12^ versus MDCK^WT^(wild-type) competition. **(B)** Cumulative mitosis events-per-cell comparing pure population induced Ras^V12^ versus competitive Ras^V12^. **(C)** Pure population cumulative mitosis events-per-cell in wild-type versus co-cultured wild-type in Ras^V12^ competition. **(D)** Cumulative elimination curves for Ras^V12^ versus wild-type competition. **(E)** Cumulative elimination events-per-cell comparing pure population induced Ras^V12^ versus competitive Ras^V12^. **(F)** Pure population cumulative elimination events-per-cell in wild-type versus co-cultured wild-type in Ras^V12^ competition. **(G)** Cumulative mitosis curves for Scr^KD^ versus wild-type competition. **(H)** Cumulative mitosis events-per-cell comparing pure population induced Scr^KD^ versus competitive Scr^KD^. **(I)** Pure population cumulative mitosis events-per-cell in wild-type versus co-cultured wild-type in Scr^KD^ competition. **(J)** Cumulative elimination curves for Scr^KD^ versus wild-type competition. **(K)** Cumulative elimination events-per-cell comparing pure population induced Scr^KD^ versus competitive Scr^KD^. **(L)** Pure population cumulative elimination events-per-cell in wild-type versus co-cultured wild-type in Scr^KD^ competition.

**Fig. 4B** displays the difference between the incidences of mitoses between competitive Ras^V12^ cells and non-competitive Ras^V12^ cells. In the competitive scenario, there is a clear reduction in mitosis count over time when compared to the non-competitive scenario, showing the context dependent behaviour of Ras^V12^ cells in competitive environments. This is further established by the equivalent graph (**Fig. 4C**) for wild-type mitosis rates in both competition and non-competition, where the number of divisions is again reduced in the competitive case.

**Fig. 4B** and **Fig. 4C** further highlights the context dependency of the behaviour of both the wild-type and mutant population. The reduction in competitive mitoses suggest that mitosis plays an integral role in the outcome of the competition for both wild-type and Ras^V12^ populations. In this context, rather than an aggressive up-regulation of the number of mitosis, as one might expect a “winner” population to engage in, there is a down-regulation making it appear as if mitoses are conserved during competition. It could therefore be considered that the conservation of mitosis from a population-level perspective is actually an indication of an organised competitive phenomena that implicates mitosis as a key part of it. To fully investigate this, a single-cell method is needed to quantify where and when these competitive mitoses are taking place.

Interestingly, **Fig. 4H** and **Fig. 4I** show a strikingly different trend to the equivalent Ras^V12^ graphs. Here, the incidences of competitive mitoses are greater than their non-competitive counterparts in both the Scr^KD^ and wild-type populations. Whereas the Ras^V12^ competition resulted in a reduction in proliferative capacity for both Ras^V12^ and wild-type during competition, the Scr^KD^ scenario seems to suggest an increase in proliferative capacity during competition. This would mean that both wild-type and Scr^KD^ populations are engaging in a form of competition that is encouraging rapid growth, rather than suppressing it. Further to this, comparing the competition mitosis counts between **Fig. 4H** and **Fig. 4I**, one can see that the rate of Scr^KD^ mitosis is almost 4-fold less than the wild-type mitoses, in what is likely a result of Scr^KD^ cell’s previously reported hyper-sensitivity to compaction [12, 14, 15]. Even though the mitotic trends are inverted when comparing the Ras^V12^ and Scr^KD^ graphs in **Fig. 4B, C, H** and **I**, the relative increase in competitive Scr^KD^ and wild-type mitoses when compared to the non-competitive mitoses is a further indication of the context dependency of this behaviour, highlighting that mitoses play an integral role in this competition.

The corresponding graphs for the incidences of elimination events (extrusion for Ras^V12^, apoptosis for Scr^KD^) are shown in **Fig. 4D-F, J-I**. The cumulative elimination counts for Ras^V12^ and wild-type cells in competition are shown in **Fig. 4D**. The number of wild-type eliminations is far greater than the number of Ras^V12^ eliminations, increasing dramatically towards the end of the time-lapse acquisitions before leveling off. This greater incidence of wild-type elimination explains the reduction in wild-type population numbers when comparing Ras^V12^ competition against Scr^KD^ competition, seen in **Fig. 3B** and **G**. This wild-type apoptotic trend is proven to be a context-dependent competitive effect in the following graph, **Fig. 4F**, where it is starkly different to the non-competitive scenario. This gives the impression that the wild-type population is facing a significant challenge in this form of competition with it’s final winner status not coming without the incidence of wild-type cells dying at an increased rate. However, it’s important to note that this increased rate of wild-type apoptoses does not determine the outcome of the competition, as the increase in the number of mitoses coupled with a similar trend of increased Ras^V12^ cell elimination, (**Fig. 4E**), is sufficient to drive the overall wild-type population count higher than the Ras^V12^ population count.

Comparing the Ras^V12^ elimination counts to the Scr^KD^, the differences between the two distinct mechanisms of competition become apparent yet again. The nature of the Scr^KD^ competition is illustrated clearly in **Fig. 4J** as relying on far greater occurrences of Scr^KD^ eliminations than wild-type. This, coupled with the far greater incidence of wild-type mitoses shown in **Fig. 4G**, points to an approximate idea of the mechanism of this competition: increased wild-type proliferation against increased incidence of Scr^KD^ elimination. Focusing on the comparison between competitive and non-competitive elimination curves in **Fig. 4K** and **L**, there is not a significant difference in the wild-type case and only a slight increase in the competitive Scr^KD^ case. This indicates that the mechanisms of this form of competition are far more subtle and diffuse than the Ras^V12^ counterparts, relying on a combination of weakness in the Scr^KD^ cell population and increased proliferation in the wild-type population. However, what is unknown is whether the increased incidence of Scr^KD^ apoptosis is cause or consequence of a concurrent increase in wild-type mitosis. Similarly, it is unknown if the increased incidence of Ras^V12^ extrusions in competitive scenarios, (**Fig. 4E**), is linked to the suppression seen in mitoses counts in the competitive wild-type populations (**Fig. 4C**).

This initial population-level analysis has revealed a striking difference between the two cellular systems, first demonstrating that competition is definitely taking place and secondly that the individual cell populations are acting in significantly differing ways to achieve the competitive outcome of mutant elimination. What is left unanswered is the question of event causality in both competitive systems. It is possible that, in the case of the Ras^V12^ population, the reduction of wild-type growth in competition (**Fig. 3E**) is due to mitosis influencing, and eventually causing, the increase in Ras^V12^ elimination (**Fig. 4E**) and subsequent outcome of Ras^V12^ competitive elimination. In the Scr^KD^ versus MDCK^WT^competition, the Scr^KD^ cells, which are far less proliferative than the Ras^V12^ cells (**Fig. 3I**), could be allowing the wildtype population to proliferate instead (**Fig. 4I**) meaning that increased growth is a consequence of the Scr^KD^ population dynamics. In order to answer this question of event causality, the incidences of both proliferative mitotic events and mutant eliminations need to be accurately quantified in relation to one another in both space and time. To achieve this, we turned to a spatiotemporal K-function analysis of the clustering of wild-type mitoses around Ras^V12^ extrusions and Scr^KD^ apoptoses.

### 2.2 K-function event clustering analysis quantitatively characterises the spatiotemporal signatures of cell competition

#### 2.2.1 Spatiotemporal counting of wild-type mitotic events around mutant cell eliminations

To investigate correlations between key events in competitive co-cultures a rigorous quantitative approach is developed that focuses on statistically analysing the distribution of mitoses around mutant cell eliminations.

Letting *x*_*i*_ be a single cell detection in the set of *n* detections (𝒟) in a time-lapse movie:

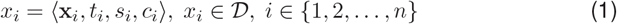

where **x**_*i*_ is the cell centroid (**x**_*i*_ ∈ ℝ^2^), *t*_*i*_ is the timestamp 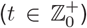, *s*_*i*_ is the state of the cell (*s*_*i*_ ∈ {interphase, mitosis, apoptosis, extrusion}) and *c*_*i*_ is the cell type (*c*_*i*_ ∈ {winner, loser}). From this set of detections, we can define subsets that correspond to mitotic (𝒟_mitotic_ ⊂ 𝒟), only winner cells (𝒟_winner_ ⊂ 𝒟), or the loser cells that contain elimination events (𝒟_elimination_ ⊂ 𝒟) and so forth. Within the dataset, we also have tracks, which correspond to a set of detections representing the trajectory of an individual cell over time:

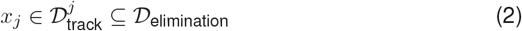

where 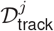 is the set of detections corresponding to track *j* containing an elimination event *x*_*j*_. Importantly, we can define the subset of all detections that are within a spatial (*r*) and temporal bin (Δ) relative to this event (*x*_*j*_):

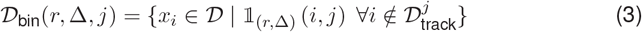

where an indicator function, as shown in equation 4, counts the number of detections within a given spatiotemporal bin (as shown in **Fig. 5**):

**Fig. 5.**
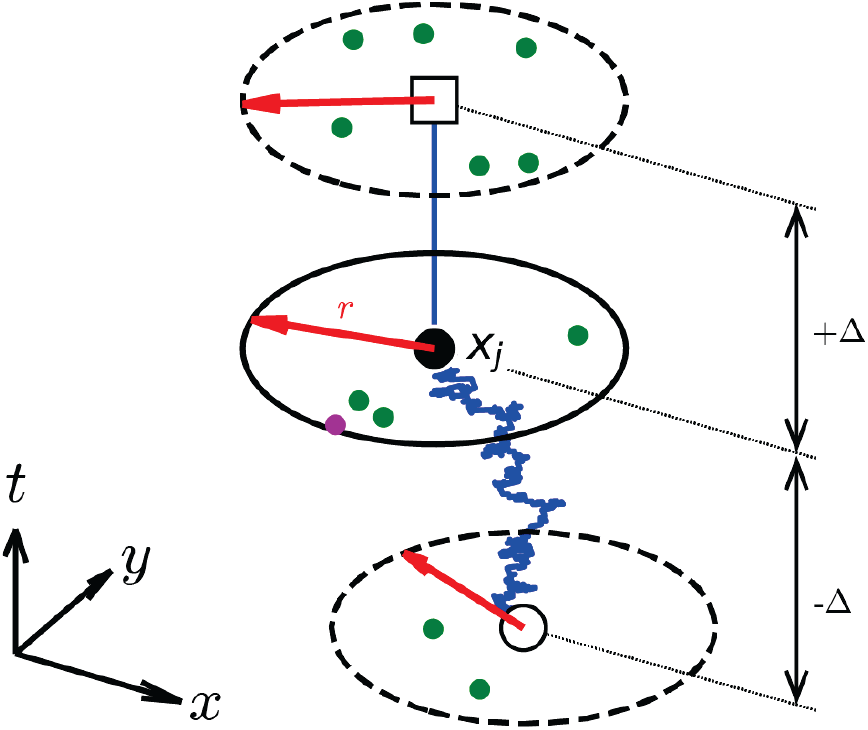
A mathematical framework for quantifying the spatiotemporal trajectory of a single elimination event (*x*_*j*_). The radius of the bin (*r*) is highlighted in red and the temporal window (Δ). Before the event we centre the window on the moving cell (blue line), whilst after the cell is eliminated, we centre the window on the last known position. Winner and loser cells are shown as green and magenta circles respectively.

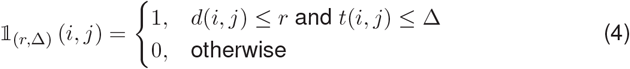

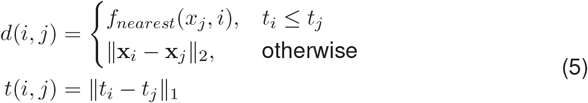

The functions *d*(*i, j*) and *t*(*i, j*) are used to define the spatial and temporal distances of each detection (*x*_*i*_) from the event (*x*_*j*_). For detections that occur before the elimination event (*t*_*i*_ ≤ *t*_*j*_), we measure the spatial distance according to the position of the cell (*f*_*nearest*_). After the elimination event, we measure the distance to the last known position, as shown in equation 5 and **Fig. 5**. Finally, we can define the set of mitotic winner cells within a spatial and temporal window of a loser cell elimination (*x*_*j*_) as the intersection between these sets:

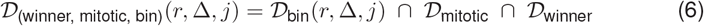

#### 2.2.2 Space-time K-function analysis to identify clusters of mitotic events

To determine whether there are specific patterns of mitotic behavior in the wild-type population that influence the elimination of mutant cells this analysis needs to consider both spatial and temporal clustering of these events. Ripley’s K-functions [24] (referred to as K-functions) are descriptive statistics used to detect and quantify clustering or dispersion in point pattern data sets. Here, we have adapted the standard K-function formulation to account for data that clusters in space and time.

A typical K-function plot features a 1-dimensional distribution showing how a point pattern data set is distributed across space. For the single-cell, spatiotemporally-resolved cell state classification data, the K-function would be 2-dimensional, showing how the number of expected mitotic events changes over space and time relative to a mutant cell elimination event.

The first step in defining a space-time K-function is to establish the spatiotemporal domain over which to conduct the analysis. In this context, this was the area, *a*(*R*), of the imaging field-of-view region *R* and the time duration of the time lapse acquisitions, *t*_*max*_ − *t*_*min*_. Next, a space-time sample estimate (*λ*_*st*_), shown in equation 7, is made, which represents the rate or frequency of events over the defined domain:

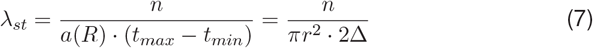

The K-function depends primarily on the indicator function, as shown in equation 4, that counts the number of events within a given spatiotemporal bin. Thus, using equations 6 and 7 we can define the space-time K-function:

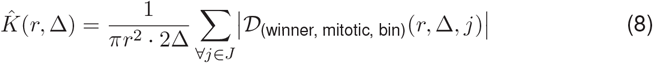

where *J* is the set of eliminated target cells (𝒟_elimination_ ∩ 𝒟_loser_) and |·| denotes the cardinality of the set. We use a range of different spatial and temporal values (0 < *r* <100 μm; −10 < Δ < 10 hours), to generate a 2-dimensional K-function plot that shows the distribution of the expected number of mitoses around a focal loser cell elimination event.

#### 2.2.3 Temporal indistinguishability test identifies statistically significant clusters of mitoses

To ascertain whether patterns of event clustering are present in the space-time K-function analysis, a temporal indistinguishability test is conducted. This involves comparing the K-function distribution of expected events, as calculated in equation 8, to a set of randomly distributed psuedo-events. These random psuedo-events are generated by shuffling the time coordinates of the observed mitotic events. This process is then repeated *N* times, resulting in *N* permutations of random pseudo-events, each one representing a null hypothesis where there is no expected spatiotemporal clustering of events. A permuted K-function, 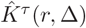 is calculated for each of the *N* set of the null-hypotheses.

The temporal indistinguishability test counts the number of instances the observed K-function, 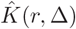 has a greater value for a given space-time coordinate (*r*, Δ) than the permuted K-function 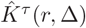 This is a simple comparison of each spatiotemporal bin in both the observed and null hypothesis 2-dimensional K-function plots. This comparison is repeated for every spatiotemporal bin and all the *N* null hypothesis K-functions, as defined in equation 9:

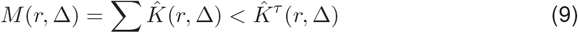

The resulting set of *M* (*r*, Δ) values are an indication of whether the observed K-function exhibits fewer event instances for a given ⟨*r*, Δ⟩ than the cumulative null hypotheses. If a large number of null hypotheses are generated, say *N* = 1000, then *M* (*r*, Δ) represents a quantitative statistical assessment of how rare the distribution of observed events are. Equation 10 is used to calculate a *p*-value for each spatiotemporal bin.

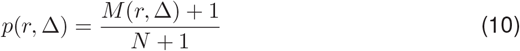

This results in a 2-dimensional, space-time plot indicating which spatiotemporal regions, relative to the focal elimination event, have statistically significant *p*-values of below 0.01. This then indicating that mitotic clustering was present in those regions. To generate a statistically significant p-value for a given spatiotemporal bin at *r*, Δ, the majority of null hypothesis K-function values must be less than the observed K-function value. This means that more events are witnessed for (*r*, Δ) in the observed K-function over the randomly generated set of events, resulting in a statistically confident assessment of event clustering.

### 2.3 K-function clustering analysis reveals a localised concentration of wild-type mitotic activity prior to- and post-Ras^V12^ extrusion

The clustering of wild-type mitoses around the apical extrusion of Ras^V12^ cells is quantified in the K-function analysis shown in **Fig. 6**. For the spatiotemporal bins that represent the local cellular environment (*<*25μm from the Ras^V12^ extrusion) in the time preceding the extrusion (−5 hours to -2 hours) there exists a concentrated pattern of p-values less than 10^−2^ showing that wild-type mitoses are clustering in this time and location. The spatial concentration of this pattern suggests that the wild-type cells are engaging in a context-dependent behaviour around the Ras^V12^ cells, with elimination being expedited by an increase in mitotic density in the hours prior to extrusion. The clustering ceases to exist from -2 hours to the extrusion onset suggesting that the process of extrusion has already been detected in the wild-type local population therefore leading to a cessation of their mitotic behaviour.

**Fig. 6.**
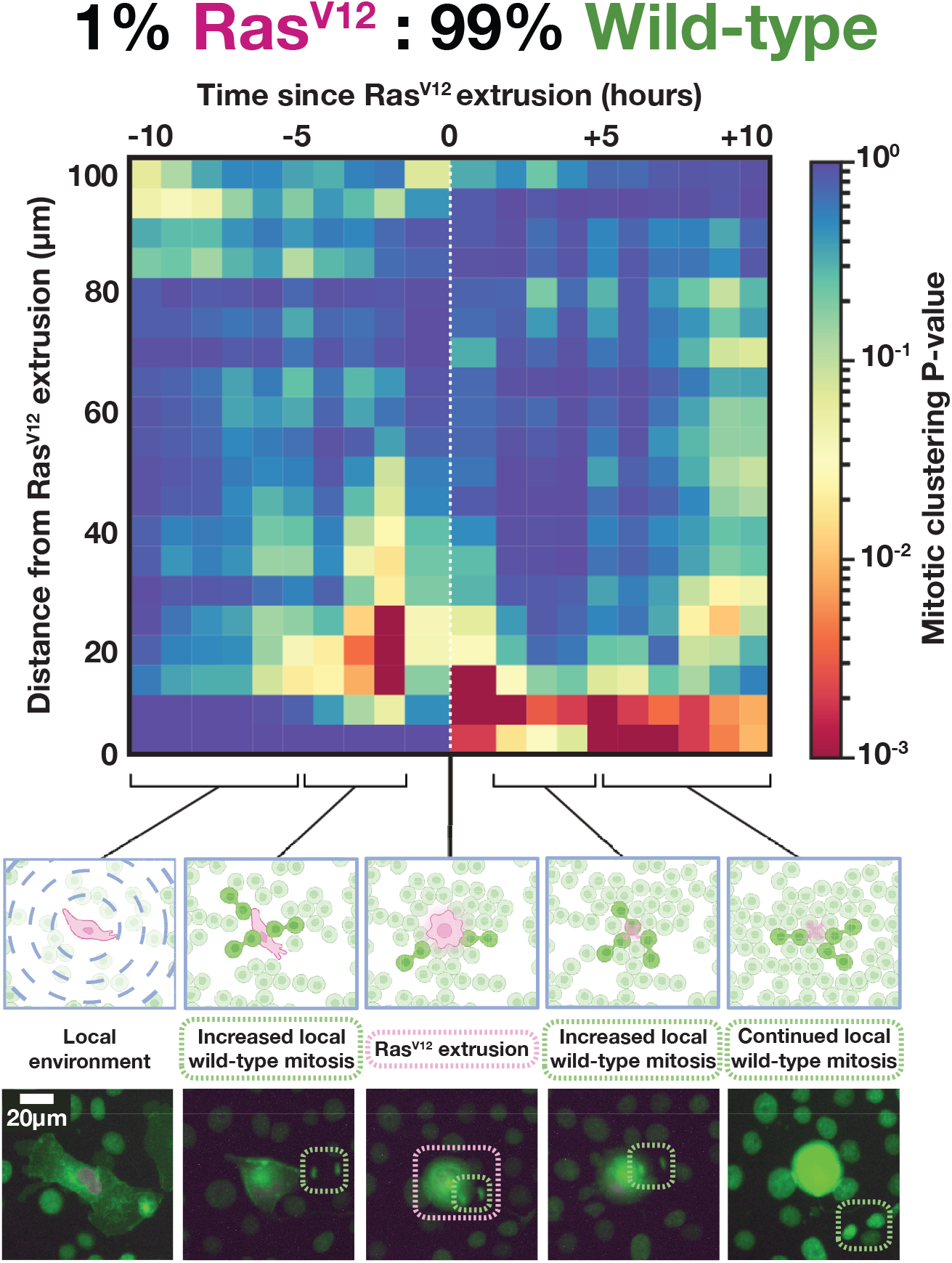
K-function clustering analysis reveals a localised concentration of wild-type mitotic activity prior to and post Ras^V12^ extrusion. A schematic representation of the biological phenomena and associated spatial characterisation of events is illustrated underneath the K-function plot. Below this representative time-lapse images are displayed, showing a single Ras^V12^ extrusion event and the reported mitotic MDCK^WT^clustering around it. [30]

This pattern of mitotic clustering continues immediately afterwards (10 hours post-extrusion) at a closer and more concentrated radial distance (*<*15μm) from the Ras^V12^ cell elimination. This could be explained by the vacation of the site once occupied by the centrally extruded Ras^V12^ cell, leaving free-space for wild-type cells to migrate and proliferate into. The evidence for this proliferation, in the form of a continued incidence of p-values less than 10^−2^, continues for many hours after the focal extrusions before tapering out approximately 8 hours post-extrusion. This suggests that alongside a causative influence, wild-type proliferative behaviour could be reacting to a decrease in local density in a similar way wild-type cells do in the context of Scr^KD^ elimination.

### 2.4 Scr^KD^ apoptosis appears unrelated to immediate, localised wild-type mitotic activity, only yielding space for diffuse wild-type proliferation post elimination

The clustering of wild-type mitoses around the apoptotic elimination of Scr^KD^ cells is quantified in the K-function analysis shown in **Fig. 7**. In contrast to the previous K-function analysis for Ras^V12^ versus wild-type, there is little evidence for localised wild-type mitotic clustering preceding Scr^KD^ apoptosis. Instead, a diffuse pattern of p-values less than 10^−2^ can be seen at further distances, representing the known up-regulation of wild-type division in the presence of mutant Scr^KD^ cells [15]. At 7 hours post-elimination a spatially concentrated pattern of p-values less than 10^−2^ (existing *<*20μm and *>*5μm away) indicates a clustering of wild-type mitoses in this region, alongside evidence for wild-type mitotic clustering many cell radii away from the central Scr^KD^ apoptosis (60-70μm away).

**Fig. 7.**
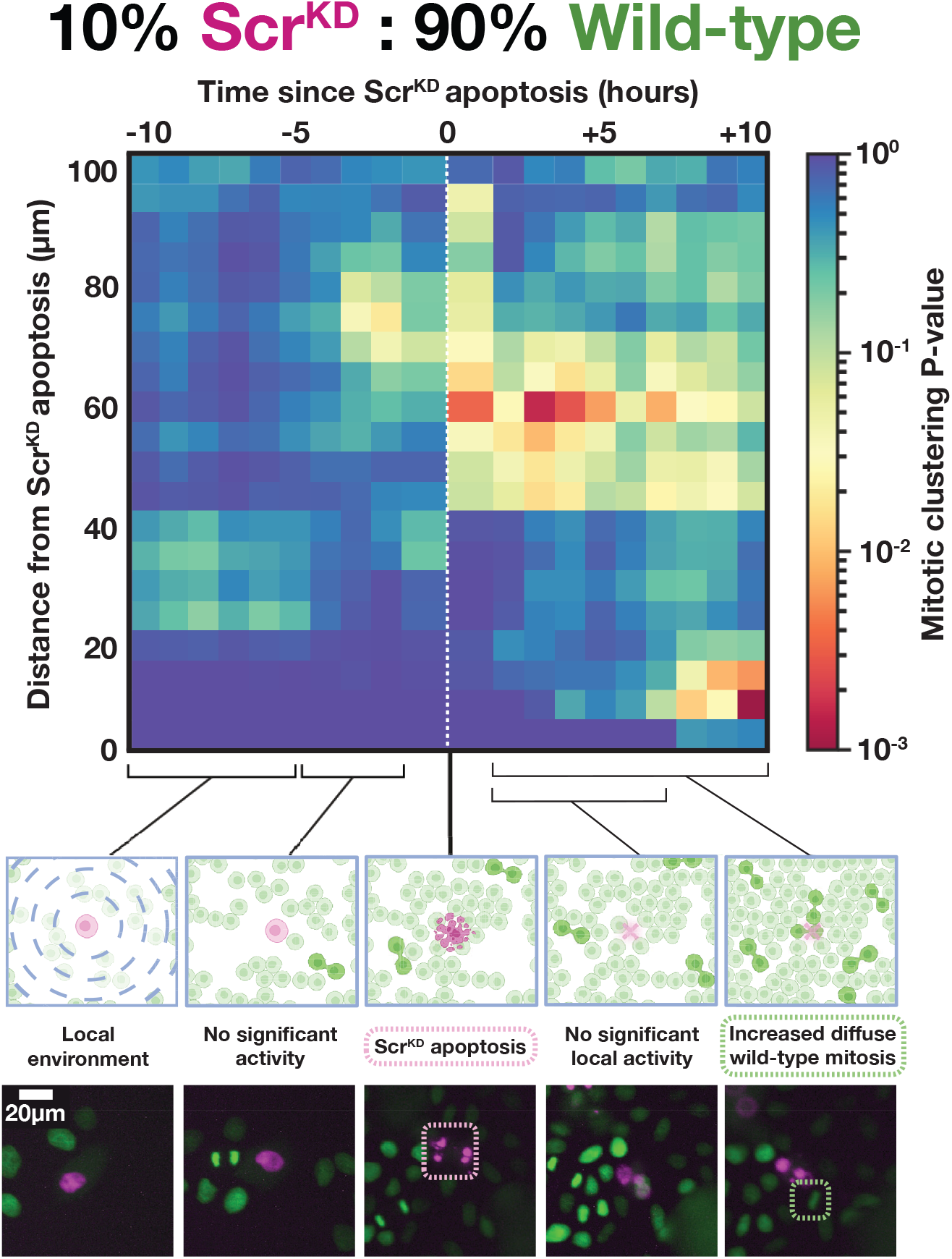
K-function clustering analysis reveals that Scr^KD^ elimination seems unrelated to immediate wild-type mitotic activity, only yielding space for diffuse wild-type proliferation post elimination. A schematic representation of the biological phenomena and associated spatial characterisation of events is illustrated underneath the K-function plot. Below this representative time-lapse images are displayed, showing a single Scr^KD^ apoptotic event and the reported lack of mitotic MDCK^WT^clustering until post elimination. Although mitosis events do occur pre-elimination, as displayed in panel two, no significant clustering is seen. [30]

This is suggestive of a space-filling effect, once the apoptotic Scr^KD^ cells are apically extruded from the monolayer post apoptosis [13], the neighbouring wild-type cells are able to repopulate the vacated space and local environment via a concentrated clustering of mitotic activity This post-apoptotic reconfiguration would be accompanied by a drop in local density, which leads to a density-dependent increase in mitotic proliferation that diffuses across the different spatial regions in the local cellular environment, according to the inverse relationship between probability of division and local density [15]. This diffusive effect is illustrated by the fact that the pattern of p-values are not spatially constrained, occurring over almost the entire 100μm range post-apoptosis.

## 3 Discussion

The aim of this study was to determine spatiotemporal signatures of cell competition for two mechanisms canonically defined as either biochemical (Ras^V12^ versus wild-type) or mechanical (Scr^KD^ versus wild-type).

Growth, mitosis and elimination dynamics of both pure and competitive co-cultures were quantified on a population-scale and context-dependent regulation of mitotic behavior in wild-type, Scr^KD^ and Ras^V12^ populations was revealed. To understand the questions raised by the population-level analysis relating to event causality, a novel K-function cluster analysis method was conducted. A strong and highly localised clustering of wild-type mitosis prior to and immediately after Ras^V12^ extrusion was seen, suggesting the wild-type population are both influencing and reacting to the ejection of Ras^V12^ mutants from the monolayer. In accordance with the novel theoretical framework introduced earlier and summarised in **Fig. 8** this suggests that Ras^V12^ vs wild-type competition can be quantified as a causal-localised and induced-localised form of cell competition. Here causal is defined as mutant cells that are influenced by actions from competitors occurring prior to cell elimination, localised is defined as events occurring approximately 0 to 40μm from the mutant cell elimination, and induced is defined as competitors reacting only once mutant cell elimination has occurred.

**Fig. 8.**
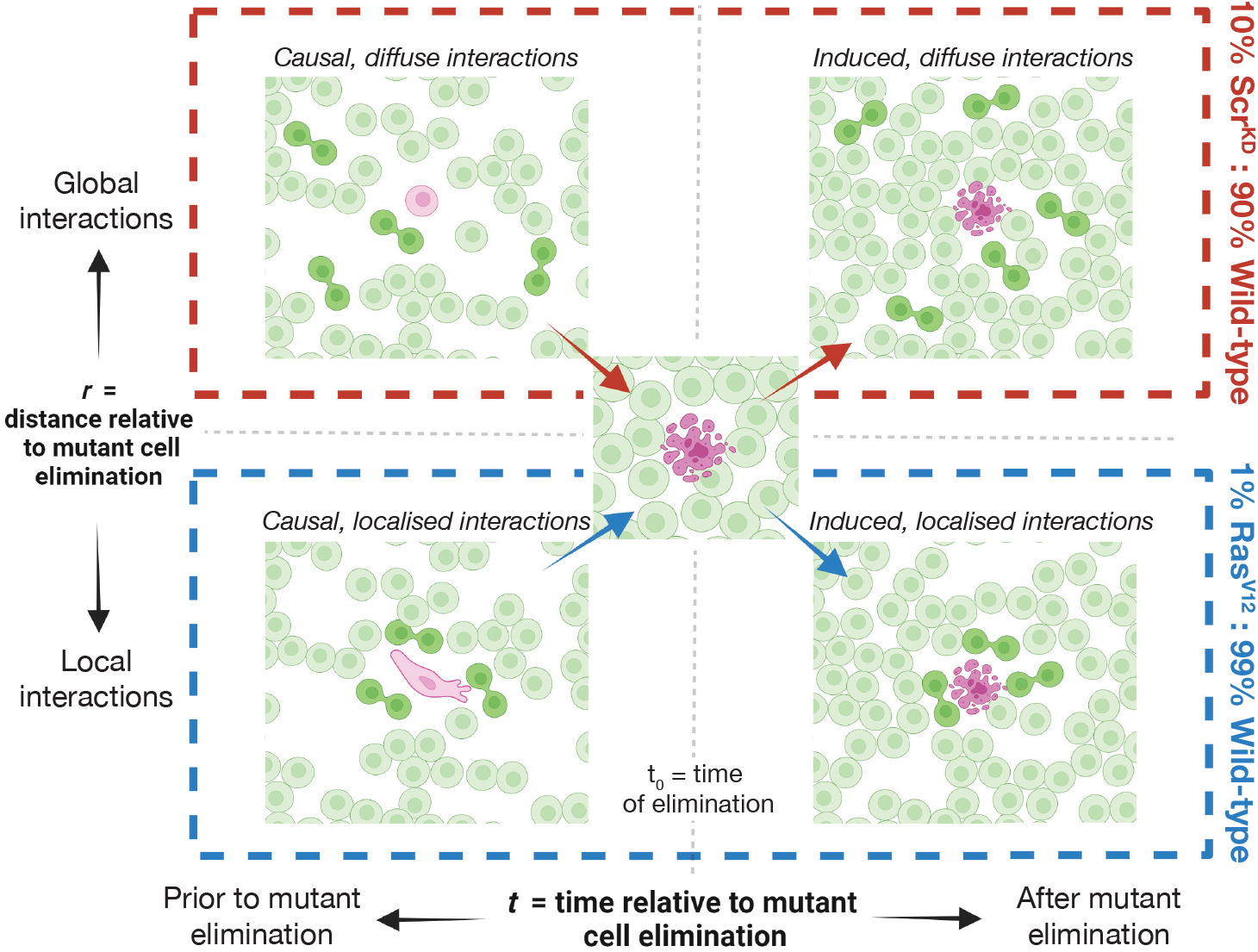
A theoretical framework for categorising different types of cell competition based on the spatiotemporal localisation of single-cell behaviors around a mutant cell elimination event. This framework yields four distinct categories, separated by which spatio-temporal dimension they are measured in. Causal interactions: mutant cells are influenced by competitor actions that occur prior to cell elimination. Induced reactions: competitors change their behaviour only after mutant cell elimination has occurred. Local interactions: key competitive events occur in the local cellular neighbourhood, defined here as approximately 0 to 40μm from mutant cell elimination. Diffuse interactions: widespread events occurring at multiple distances relative to the mutant cell but that still have an influencing effect on the outcome of the competition. [30]

K-function cluster analysis additionally revealed that following a Scr^KD^ apoptosis there is both a strong, delayed cluster of localised wild-type mitosis suggesting a reactive space-filling mechanism, and a broader prevalence of mitosis events across the monolayer. As shown previously in [15] and **Fig.4** wild-type cells up-regulate their growth in the presence of Scr^KD^ cells. This is reflected in **Fig. 7** as the mild increase in clustering of mitosis events *<*40μm away prior to the Scr^KD^ apoptoses. This places Scr^KD^ competition as possessing causal and induced characteristics, both occurring as a more diffused pattern of behaviour than observed in the Ras^V12^ scenario. In the framework described in **Fig.8** diffuse interactions are defined as events that are widespread, occurring at multiple distances relative to the mutant cell, and have an influencing effect on the outcome of the competition.

Taken together, these analyses clearly differentiate two distinct reactions of wild-type populations to the threat of a mutagenic cellular invasion. These results suggest that more nuance is required when classifying types of competition and that relying on clearly quantifiable metrics, such as the spatiotemporal localisation of key competitive events, is a more suitable approach than assigning a binary classification of biochemical or mechanical, especially considering that all cellular processes are inherently biochemical or mechanical in varying capacities.

Our method of characterising these signatures of competition focused on explicitly quantitative measures of the spatial and temporal localisation of three sets of key competitive events: elimination, apoptosis and extrusions for wild-type, Scr^KD^ and Ras^V12^ populations. These events are key because they represent the single-cell origins of the two outcomes of cell competition: proliferation versus elimination. By tracking the dynamic, single cell level processes that influence the emergent phenomena of cell competition we can gain a deeper understanding of the rules that govern these phenomena. Our approach has been designed and implemented as a template for further classification endeavours within the realm of cell competition and is applicable to many other cell systems that are currently not characterised, as well as more general biological models outside cell competition, such as in the field of development and cell differentiation.

Further work is needed to confidently assert causative effects between competitive events and the outcome of cell competition. Future experiments to address this could involve inhibiting mitosis in the wild-type population through a live-cell perturbation assay and subsequently recording the frequency of mutant elimination. If the frequency of Ras^V12^ extrusions decreases then this would serve as further confirmation of the competitive influence of wild-type mitoses in this system. If the frequency remains the same then the role of other cellular behaviours should be investigated, such as concerted migration of cells, concentrated regions of cellular density, and co-ordinated cellular signaling mechanisms. These behaviours all represent more subtle, diffusive mechanisms of competition that may be present in the absence of preemptive clusters of mitoses.

Our study provides a comparative assessment of two canonically defined forms of cell competition, offering a nuanced characterisation of the single-cell mechanisms driving mutant elimination from a spatiotemporal and statistical perspective. These results prompt a reevaluation of the prevailing definitions of each competition type. We propose a shift towards a more mechanistic understanding of cell competition, potentially reshaping our approach to studying these intricate and dynamic cellular processes using cutting-edge, robust quantitative methods.

## 4 Materials and methods

### 4.1 Cell Line Preparation

The MDCK wild-type cells used in this project were a gift from Prof. Yasuyuki Fujita (University of Kyoto, Japan). The Ras^V12^ cell line is described in Hogan et al. [9]. The Scr^KD^ cell line is described in Norman et al. [13]. Stably fluorescent tagged histone markers, H2B-GFP for the wild-type population and H2B-RFP for the Ras^V12^ and Scr^KD^ mutant populations, were developed to differentiate competitive cell types and to track individual cell cycle progression. The Ras^V12^ population also possessed a Ras^V12^ -GFP marker to indicate cytoplasmic Ras^V12^ distribution. The Scr^KD^ population possessed a GFP marker to indicate cytoplasmic cell shape.

### 4.2 Cell Line Maintenance

Cells were grown in DMEM culture media (Thermo-Fisher) supplemented with 10% tetracycline-free FBS (Clontech, 631106), 1% sodium pyruvate (NaPy, Sigma-Aldrich) and 1% penicillin streptomycin. Cells were grown to 80% confluency in a humidified incubator (37°C, 5% CO_2_) to allow for the development of epithelial functionality and cell-to-cell adhesion before being passaged into new T-25 flasks.

Cell passaging was performed by firstly aspirating the old media from the monolayer, then rinsing the cells with double the volume of PBS, before adding 1mL of trypsin-EDTA solution (0.05% trypsin, 0.53mM EDTA) and incubating for 20 minutes. After verifying that the cells had fully detached, the trypsin-cell solution was diluted by adding 4mL of warmed DMEM in the T-25 flasks and agitating the solution using a pipette to fully dissociate any remaining clusters of cells. 5 mL of fresh DMEM was then added to a new T-25 flask followed by 350μL-500μL of suspended cell solution.

Cells were tested monthly for mycoplasma (MycoAlert Plus Detection Kit, Lonza, LT07-710) and if an infection was found the cells were treated with 1/1000 concentration puromycin for 2 weeks until the infection was eliminated and the sample tested negative.

### 4.3 Induction of Mutation

Both the Scr^KD^ and Ras^V12^ cell lines were cultured as wild-type up until the induction of the shRNA system to express the mutation of interest for an experiment. Scr^KD^ cells were induced with 1μg/mL doxycycline (Sigma-Aldrich, D9891) for 70 hours before seeding in preparation of an imaging sample. Ras^V12^ cells were induced directly in the imaging well at a concentration of 1μg/mL doxycycline.

### 4.4 Preparation of Pure Population and Competition Assays

Cells were seeded in 24-well imaging plates (Ibidi) at a density of 1×10^−3^ cells/μm^2^. Pure population wild-type, Ras^V12^ and Scr^KD^ were seeded at densities of 3×10^−3^ cells/μm^2^, 1×10^−3^ cells/μm^2^, and 3×10^−3^ cells/μm^2^ respectively. Competition assays of wild-type and Ras^V12^ were mixed at 95:5, 97:3 and 99:1 ratios and seeded at a density of 3×10^−3^ cells/μm^2^. Competition assays of wild-type and Scr^KD^ cells were mixed at 90:10 ratios and seeded in 24-well imaging plates (Ibidi) at a density of 1×10^−3^ cells/μm^2^. After seeding, cells were incubated for 3 hours to form basolateral adhesions before starting an image acquisition, and were typically imaged for 4-7 days, dependent on the competition assay and experiment-specific observations.

### 4.5 Widefield Fluorescence Microscopy

Time-lapse imaging was performed on a bespoke automated epifluorescence microscope built inside a standard CO_2_ incubator (Thermo Scientific Heraeus BL20) operating at 37°C, 5% CO_2_. The brightfield illumination uses a fibre-coupled green LED (Thorlabs M520L3, 530 nm). Widefield epifluorescence illumination was provided by a LED light engine (Bluebox Optics Niji). The LEDs are combined using a dichoric beamsplitter (Semrock) and focused onto a 20x air objective (Olympus Plan Fluorite, 0.5 NA, 2.1mm WD). The sample location was controlled by high performance encoded motorised XY and focus motor stages (Prior H117E2IX, FB203E and ProScan III controller). Images were acquired with a 9.1MP CCD camera (Point Grey GS3-U3-91S6M). Cameras and light sources were synchronised using TTL pulses from an external D/A converter (Data Translation DT9834) to ensure minimal light exposure. The sample was kept at the correct humidity using a custom built humidification chamber fitted with a thermocouple and humidity sensor.

### 4.6 Confocal Microscopy

Volumetric images of fixed Ras^V12^ and Scr^KD^ competitive scenarios were acquired using a 100x oil immersion objective (1.40 N.A., Olympus) mounted on an Olympus IX83 inverted microscope equipped, with a scanning laser confocal head (Olympus FV-1200). Images were acquired with a *z*-step of 1μm and total field of view of 211.97μm x 211.97μm. Imaging samples were excited at 488 nm and 562 nm for the green and red fluorescence profiles, respectively.

### 4.7 Single-cell segmentation and tracking

Nuclear segmentation of the widefield fluorescence images was performed using *Cellpose*[18], *StarDist* [19, 20] or a residual U-Net [21]. Cell mitotic state labelling was then applied using a dedicated convolutional neural network as previously described [15, 22, 23]. Finally, each cell instance was tracked over time using our *btrack* [15, 22] package prior to manually verifying key competitive events in the mutant populations using a custom plugin for napari [25]. All image processing was performed on a Dell Precision workstation running Ubuntu 20.04.03 LTS with 32GB RAM and an NVIDIA GTX1080 GPU.

### 4.8 Identification of key cellular events involved in competition

Elimination events in the two competitive systems take the form of Scr^KD^ apoptosis, MDCK^WT^apoptosis, Ras^V12^ apical extrusion or MDCK^WT^apical extrusion. To investigate any casual link between these events and wild-type mitoses on a cellular scale, we first needed to annotate their occurrence in the microscopy data in space and time.

Although there are automated methods for event detection [22, 26, 27, 28], we chose to manually identify all of our key elimination events to ensure that our analysis was based on a well-defined ground truth set of events. This approach also had the benefit of creating a corpus of ground-truth data relating to these single-cell events to inform future automated classification approaches. Due to the much greater number of division events taking place, a previously established automated approach was used [15].

#### 4.8.1 Automated identification of MDCK^WT^, Ras^V12^ and Scr^KD^ mitoses

Mitosis represents a key competitive event in the wild-type cellular populations. In contrast to the manually identified elimination events, wild-type mitoses were identified using a previously described automated method [15, 22, 23]. Observation of fluorescent H2B-GFP condensation of pro/meta-phase and the clear separation of chromatid pairs during anaphase allows three different mitotic classifications: prophase, metaphase, and anaphase; as well as interphase. Errors in mitotic classification were corrected using a temporal model of the cell cycle [29] implemented as a hidden Markov model (HMM), as detailed in [15]. This resulted in any tracks that contained a metaphase to anaphase transition being reliably classified as a mitotic event without need for further manual verification. The total number of automatically identified wild-type mitoses was *N*_WT mitosis_ = 3399 across *N* = 20 Ras^V12^ vs wild-type competition time lapse acquisitions, *N*_WT mitosis_ = 11, 520 across *N* = 18 Scr^KD^ vs wild-type competition acquisitions, and *N*_WT mitosis_ = 11, 486 across *N* = 17 pure population wild-type acquisitions. The total number of automatically identified Ras^V12^ mitoses was *N*_Ras mitosis_ = 143 across *N* = 20 Ras^V12^ vs wild-type competition acquisitions and *N*_Ras mitosis_ = 2126 across *N* = 5 pure population Ras^V12^ acquisitions. The total number of automatically identified Scr^KD^ mitoses was *N*_Scr mitosis_ = 192 across *N* = 18 Scr^KD^ vs wild-type competition acquisitions and *N*_Scr mitosis_ = 241 across *N* = 10 pure population Scr^KD^ acquisitions.

#### 4.8.2 Manual identification of Ras^V12^ and MDCK^WT^cell apical extrusions

In contrast to the well-defined morphological changes and relatively short timescale of mitosis, an extrusion event can occur over many hours and may not have as clearly defined or sudden morphology change. Therefore, we manually identified extrusion events in the dataset.

**Fig. 2A** illustrates a typical morphology change of a single Ras^V12^ cell undergoing an apical extrusion from the monolayer as imaged with the widefield fluorescence microscope. The time point of extrusion was defined as the point at which the cell has rounded up and moved out of the focal plane (as seen in the illustrative image sequence in **Fig. 2A**). This extrusion was confirmed in every instance by following the cell for the rest of the time-lapse to confirm that it either underwent a further apoptosis or remained extruded and out of the focal plane, unable to reintegrate into the monolayer. To further verify this morphology change as a Ras^V12^ extrusion, several competition experiments were fixed post-time-lapse acquisition and imaged on a confocal microscope. 5 positions over 2 experiments were checked for apical extrusions to see if the expected phenotype was present and an example of this is shown in **Fig. 2C**. A total of *N*_Ras extrusion_ = 90 and *N*_WT extrusion_ = 121 were manually labelled from *N* = 20 Ras^V12^ vs wild-type competition acquisitions. A total of *N*_Ras extrusion_ = 12 were manually labelled from *N* = 5 pure population Ras^V12^ acquisitions. A total of *N*_WT extrusion_ = 25 were manually labelled from *N* = 17 pure population MDCK^WT^acquisitions.

#### 4.8.2 Automated identification of Scr^KD^ and MDCK^WT^cell apoptoses with manual time point verification

A typical phenotype of the competitive elimination of Scr^KD^ cells is shown in **Fig. 2B**. Unlike extrusion, the time range of apoptosis occurs over a matter of minutes and exhibits a sudden and recognisable morphology change due to the fragmentation of the fluorescent H2B marker present in all the cell types. Identification of apoptotic cell state transitions were already factored in as a part of the tracking pipeline previously published by Bove [15], so this automatic classification was used to bootstrap the identification process. After automatic identification, we manually verified the precise time point of each apoptosis, resulting in a total of *N*_Scr apoptosis_ = 101 and *N*_WT apoptosis_ = 121 from *N* = 18 Scr^KD^ vs wild-type competition acquisitions. A total of *N*_Scr apoptosis_ = 258 from *N* = 10 pure population Scr^KD^ acquisitions. A total of *N*_WT apoptosis_ = 23 from *N* = 17 pure population MDCK^WT^acquisitions. Where both extrusion and apoptosis events occurred for each cell population they are combined and relabelled as ‘elimination events’ for the purpose of the population-level analysis.

### 4.9 Temporal-alignment algorithm for population-level analysis

Bulk analysis of the number of single-cell measurements over time was used as a starting point to verify that cell competition was taking place and to characterise said competition on a population scale. In order to compare population dynamics over different biological and technical repeats a growth curve alignment algorithm was used to allow for a comparison of the number of cells and/or mitoses over time for different experiments. In this alignment algorithm a reference curve was chosen for each condition and the remaining growth curves were then overlaid onto the reference curve by implementing a translation in the temporal dimension (*x*) until the root mean square error is minimised. For competition experiments where there are two sets of growth curves (i.e. for wild-type and mutant cell types present in the same field of view), the wild-type cell curves are aligned using the above method, and the corresponding mutant cell curves are shifted by the same value in *x*. Once the growth curves from all experiments in each condition have been temporally aligned in this way a crop is applied to all curves in competitive and wild-type pure population conditions to ensure the initial cell numbers for the MDCK^WT^cells are consistently 100.

### 4.10. Software availability

The K-function analysis scripts can be found at https://github.com/nthndy/cell-comp-analysis. The Bayesian tracking library is available at https://github.com/quantumjot/btrack.

## 5 Author contributions

ARL and GC conceived and designed the research. ND, JM, GV and MK performed experiments. ND developed the space-time K-function method and performed K-function computational analysis. JM performed the cell population growth and event dynamics computational analysis. All authors evaluated the results and wrote the paper.

## 6 Acknowledgements

This work was supported by a MRC studentship to ND and an EPSRC studentship to JM. GV and MK were supported by BBSRC grant BB/S009329/1. We thank Y. Fujita for the kind gift of cell lines used in this work. We also thank members of the Lowe and Charras laboratories for discussions and technical support during the project. ARL acknowledges the Turing Fellowship from the Alan Turing Institute. ARL and GC acknowledge the support of BBSRC grant BB/S009329/1.

## Notes

### Competing Interest Statement

The authors have declared no competing interest.

